# Discrete Modeling Dynamical Systems That Determine the Role of Biodiversity in Different Regimes of Using Resources at Different Levels of Organization of Living Matter

**DOI:** 10.1101/281279

**Authors:** Yu.G. Bespalov

**Affiliations:** V. N. Karazin Kharkiv National University, Ukraine

**Keywords:** biodiversity, usage of resources, strategies of performance, biosafety

## Abstract

The paper aims at mathematical modeling of dynamic systems that determine the role of biodiversity in different modes of resource usage at different levels of the organization of living matter. As a measure of biodiversity, the number of different strategies of the performance of a biological system is used, which differ in ways of usage of the resources by this system. This measure can be supplemented, and in some cases, replaced by the degree of evenness of the intensity of different ways of using resources. As a result of modeling with use of a new class of mathematical models called Discrete Models of Dynamical Systems, the appearance of matrices representing the sets of these strategies corresponding to combinations of values of the system’s parameters in columns, strategies observed for the regimes of maximum and limited usage of resources was analyzed. Using the case study related to a local fish population in a pond, it was shown that, in comparison with the regimes of limited use of resources, the regime of maximum usage distinguishes by a larger degree of evenness of the intensities of different ways of using resources by the system. Different size-age groups of the population play the role of resources. A similar result was obtained earlier for different regimes of the performance of the human cardiovascular system; in this case, the heart rate and blood pressure played the role of different ways of resources usage. The similarity of the results obtained for such different biological systems allows us to hope for the prospects of developing universal system approaches to the use of generalized biodiversity parameters for the diagnosis of the state of bioobjects, in particular, with the help of aerospace methods. The necessity of such methods is significantly increasing in the context of global climate change.

## Introduction

Now, the problem biodiversity’s role in the performance of systems at different levels of organization of living matter is very topical from the point of view of theoretical biology. From a practical point of view, this topicality is determined by the need to estimate the risks of the emergence of various threats to biosafety in connection with the loss of stability by biological objects due to a violation of the mechanisms of their homeostasis. These risks significantly increase in connection to global climate changes in the context of increasing pressure of the human civilization on nature. One of the important reasons for strengthening this pressure is the need to increase the productivity of the agriculture sector of the World economy. Increasing the productivity of artificial ecosystems in plant cultivation is usually associated with measures for reducing biodiversity in them. This is due to the need to create optimal conditions for the growth of one biological species giving a certain economic effect and suppressing others species — its competitors and enemies. A similar conflict occurs in other spheres of the agriculture sector of the economy, for example, in fish farming, where in many cases measures are being taken to combat coarse fishes or enemies of feed organisms. While preserving certain technologies, low biodiversity is combined with high productivity and, in a certain sense, stability. If such technologies cannot be provided (in particular, in biological wildlife communities), low biodiversity causes the risk of a sharp decline in productivity which can be interpreted as an aspect of the loss of stability. These risks are related to the emergence of conditions unfavorable for the dominant species and the lack or insufficient abundance of species for development of which these new conditions are favorable. As a consequence, a sharp drop may occur in the overall productivity of a biological community. The increase in productivity in the form of mass development of a new dominant species is also possible. For example, toxic cyanobacteria can reveal mass development in the case of “blooming” of reservoirs, conditions in which have changed due to eutrophication. Such a change of the dominant species is generally unfavorable from the point of view of human interests and often, as in the mentioned case, generates threats to biosafety. Resonant examples of such threats are the following:

-in the Baltic Sea, where toxic cyanobacteria form “bloom patches” posing problems for recreation and fisheries [1];

-potential thread on Lake Kinneret, the main source of drinking water supply in Israel [2].

The viewpoint on biodiversity as a factor in the stabilization of communities of living organisms has a long history and now is very popular among specialists and community. But due to the lack of unambiguous universality of the factual material, it is very discussable. The concept of optimal variety, recently proposed by E.N. Bukvareva et al. [3-17], suggests trade-off views in certain specific cases. An essential aspect of this concept is consideration of the connection between an optimum value of the variety of the system and amount and availability of resources that system uses.

Along with the number of species, the evenness of amounts of criteria for their significance (biomass, abundance etc.) is used to characterize biodiversity. For numerical evaluation of both these parameters, for about half a century they use approaches based on the Shannon index [19]. It should be noticed that use of these approaches entails a decrease in the role of rare species that contradicts the established paradigm of nature protection. R. Margalef, one of the pioneers of applying the Shannon index in ecology wrote at his time that a researcher has to see the structure of relations in communities of living organisms behind diversity indicators [20]. Successes of modern mathematics and information technologies assume the great role of mathematical models as a tool for such the vision. But it should be marked the difficulties in collection of necessary factual data, not only for the formal description of this relations’ structure but even for detection of the species composition of a certain biological community at a certain location in space and time. These difficulties were quite essential even in the era when naturalists described biological systems performing under relatively unchanging, evolutionary patterns when for preventing the threats to biosafety it was only needed to minimize human intervention in the life of nature. (Nature, according to the known “law” of Commoner, “knows better”).

In conditions of global climate changes increasingly affecting the appearance of entire natural zones, these difficulties increase several times. However, today’s possibilities of studying biological systems, in particular, by remote methods are also increasing: satellite and cheaper methods involving the use of unmanned aerial vehicles (UAVs). These methods allow researchers to collect in a short time information on the status of biological objects in large, sometimes difficult-to-reach areas, which can be of paramount importance in extreme situations caused by large-scale natural and man-made disasters. But in regard to arrays of these data and their shortcomings, three important notes should be mentioned.

First, in many cases these arrays will contain information not about those parameters that are traditionally recorded during environmental studies; this implies the use of generalized indicators reflecting the relations between different parameters inherent to certain systems and their states, in particular, the relations between parameters that reflect diversity, productivity, and stability (it is assumed that certain types of these relations can have similarities for parameters having different physical and/or biological meaning).

Second, even the parameters traditionally recorded by ecologists can be measured with a lower accuracy than usually; this makes it possible to use ranked parameters and recording their values in conditional scores.

Third, such arrays can have gaps (invoked by cloudiness, other visibility-impairing circumstances, non-flying weather, etc.); these gaps will not allow obtaining a sequence of states of an observed object in real time, meanwhile, the parameters supporting the homeostasis of living systems and their stability are closely related to the nature of the dynamics of the processes occurring in them.

A new class of mathematical models developed at the V. N. Karazin Kharkiv National University, Ukraine, and already applied to study various systems [21-25] enables to obtain a formalized description of mechanisms of homeostasis and dynamics of systems of different nature. For models’ identification, this tool can use data having above mentioned drawbacks. Such models bear the generic name of Discrete Models of Dynamical Systems (DMDS). The DMDS makes it possible to describe the inter- and intra-component relationships in a system, which are caused by inter- and intra-component positive and negative effects. On the basis of the structure of these effects, an idealized trajectories of the system (ITS) reflecting the cycle of changing the system’s states can be constructed for certain initial conditions. We keep in mind the states corresponding to specified unique combinations of the values of system’s components in the cycle, which are observed in the cycle at different moments of time. In [26], it was proposed to consider a set of these combinations as a set of strategies of the system allowing it to support the mechanisms of homeostasis, which determine the type of the cycle with the named combinations on conditional steps. Each of these combination-strategies can be assessed by the evenness of values of all or a part of the components of the system, as well as by values of productivity, or by the efficiency of various aspects of its performance. These values of efficiency or productivity can be represented at different time moments of ITS by values of the system’s components modeled by the DMDS. These components can also be interpreted as different ways of using resources by the system to ensure a certain pattern of its performance (or, performance in general). Notice that in the above-mentioned concept [3-17] of optimal diversity, the criteria for optimality of biodiversity, related to the quantity and availability of resources used by the system, play an important role. It should be noted that in a number of cases the degree of evenness of the values of the system’s components can be accepted as a main measure of diversity. (For example, in cases when the number of the system’s components does not vary or the number of all components of rare occurrence cannot be taken into account).

In connection with these considerations, the problem of the study of relations between the nature of usage of resources by the system and the evenness of intensity of various methods of this usage seems meaningful.

The results of such the study conducted with use of the DMDS on the data of the human cardiovascular system are presented in [27]. In this work, the differences in the degree of evenness of values of heart rate (HR) and blood pressure (BP) for different types of performance of the body were described.

We are dealing with regimes with maximum usage or saving of resources.

These regimes hereinafter are referred to as the regime of maximum usage of resources (RMUR) and the regime of limited usage of resources (RLUR), though these names are quite conventional.

In this case, HR and BP play the role of different ways of resources usage to change the speed of blood flow. In the first case, there is a greater degree of evenness of these regimes (that is, expressed in the conditional scores of HR and BP) than in the second. In this case, such the degree of evenness is a significant aspect of biodiversity.

These results give certain grounds to suggest a working hypothesis, according to which the degree of evenness of different ways of system’s resources usage can indicate the performance of regimes of their maximum usage or saving. We keep in mind the systems at different levels of organization of living matter and different types of resources necessary for the performance of the systems.

Within the framework of this working hypothesis, the differences between these regimes in the character of relationships of this degree of evenness with the degree of dominance of specific ways of using the resource can be considered. For example, in the case of a fish population, size-age classes will be matched to these modes. The classes can be ranked, according to their size, according to intensiveness of consumption of natural forage resources per individual. Using such and other methods of ranking, appropriate approaches to solving the problem of this study may be suggested. These approaches have some features similar to bioindication methods for the purity of reservoirs and watercourses with use of methods of indicator organisms in conjunction with biodiversity criteria.

The verification of this hypothesis on the data of the system, the biological nature of which is quite different from a human body, would create certain prerequisites for development of an universal approach to the problem of the connection of the regimes of performance of systems of different nature with the evenness of various ways of using resources by such systems. In a long run, we can deal in this context with other aspects of biodiversity, along with evenness.

This work aims at such the verification with use of the DMDS on the data of a local fish population (*Karassius karassius*) in a pond.

## Material and methods

To attain the aims mentioned above, aspects of evenness of the intensiveness of the ways of using resources in different regimes of the system performance were analyzed. For this purpose, a comparative analysis of the appearance of ITSs calculated with use of the DMDS on the actual data of the literature source [27] was carried out. The data describe the character of the fish population size structure (*Karassius karassius*) in a pond before and after the catch that reduced the density of the population and improved the conditions of use of a natural forage reserve by fishes. The systemic effects obtained from the data of the fish population were compared with those described earlier in [27] for the human circulatory system mentioned above.

The ITS was calculated with use of the DMDS with the measure of proximity based on Spearman correlation and dynamics based on an extended interpretation of the von Liebig law [21].

## Results and discussion

ITSs obtained on the above-mentioned actual data reflect the dynamics and distribution of different values of the system’s components, which are different values of the number of fishes belonging to the three following dimensional classes:

having body length from 25 to 45 mm (L 25-45 mm)

having body length from 65 to 85 mm (L 65-85 mm)

having body length longer than 95 mm (L- 95 mm).

Besides the three parameters, the fourth parameter referred to hereafter as latent (LK) was added to the model. Correlations between this latent component and other components were assumed equal to zero. The biological meaning of such correlations may be different. In particular, it can consist in the lack of connection between components or the presence of the “plus-minus” relationship, which plays an important role in the mechanisms of homeostasis of systems of different nature.

Table 1 presents the ITS corresponding to the state when the catch has not yet reduced the density of the population and, as a result, has not improved the conditions for use of natural forage by fishes. That is, we keep in mind the state with greater restrictions on usage of these resources by the system. As can follow from Table 1, in this state, the coincidence of the maximum values of the number of size classes L 65-85 mm and L- 95 mm is revealed only at one (eleventh) time moment.

**Table 1.** Idealized trajectory of the system representing the dynamics of the amount of the size classes of the body length of *Karassius karassius* in the period prior the experimental population decline.

|  |  |  |  |  |  |  |  |  |  |  |  |  |  |  |
| --- | --- | --- | --- | --- | --- | --- | --- | --- | --- | --- | --- | --- | --- | --- |
| L 25-45 mm | 1 | 1 | 1 | 2 | **3** | **3** | **3** | 2 | 1 | 1 | 1 | 1 | 1 | 1 |
| L 65-85 mm | 1 | 1 | 1 | 1 | 1 | 1 | 1 | 1 | 1 | 2 | **3** | **3** | **3** | 2 |
| L>95mm | 1 | 2 | 2 | 1 | 1 | 1 | 1 | 2 | **3** | **3** | **3** | 2 | 1 | 1 |
| LK | 1 | 2 | **3** | **3** | **3** | 2 | 1 | 1 | 1 | 2 | 2 | 1 | 1 | 1 |
| Conditional time moments | 1 | 2 | 3 | 4 | 5 | 6 | 7 | 8 | 9 | 10 | 11 | 12 | 13 | 14 |

In Table 2, this coincidence presents on three (1st, 2nd, and 3d) moments. Moreover, for this table, the above mentioned three moments coincide with the maximum value of the number of individuals belonging to the specified classes. Such a relatively high coincidence frequency can be considered as a manifestation of high evenness of the number of specified size-age classes in the area of high values.

**Table 2.**
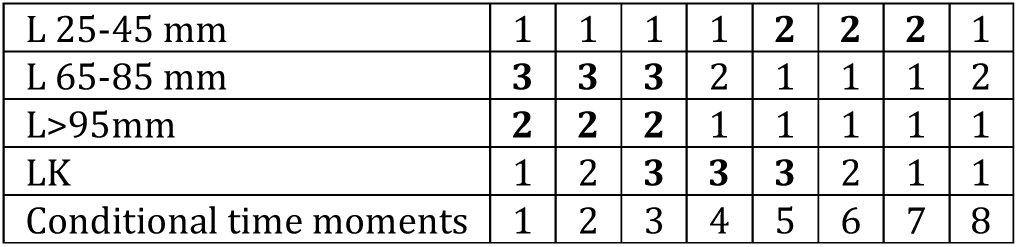
Idealized trajectory of the system, representing the dynamics of the number of size classes of the body length of *Karassius karassius* in the period after the experimental population decline. Notation as in Table 1.

We will interpret different size classes of fishes as different ways of using the resource of a natural forage reserve by a population of fishes. We also consider the number of individuals of these size classes as a measure of the intensiveness of resource’s use. Based on such assumptions, it is possible, in this case, to draw a conclusion about the connection between the evenness of the manners of resource’s usage with the regime of such the usage. Specifically, we are talking about the following. Before the moment when the catch has reduced the density of the population and, accordingly, improved the conditions for usage of natural forage by a separate individual, there had been the regime of limited usage of a resource similar to the mentioned above and described in [27] for RLUR of the circulatory system. In this case, RLUR corresponds to a relatively small degree of evenness of the number of L 65-85 mm and L> 95 mm size classes. Situations after the catch, which has reduced the density of the population and, accordingly, improved the conditions for usage of natural forage by a separate individual, corresponds to RMUR. It is obvious from the comparison of the appearance of the ITSs shown in Table 1 and Table 2, the degree of evenness of amounts of L 65-85 mm and L- 95 mm size classes in the area of their high values for RMUR is significantly higher than for RLUR. This can be, in particular, a manifestation of weakening the food competition between similar size classes after decreasing the population density. It is also worthy of note that these differences in the degree of evenness correspond to certain ways of using the resource, which makes it possible to draw an analogy with the use of indicator organisms and indicators of biodiversity in ecology.

In rows — values of the system’s components: the latent component and body lengths expressed in conditional scores (1— low, 2 — medium, 3 — high). In columns — conditional time moments. Abbreviations of the system’s components are clarified in the text. The maximum values for the table are indicated in bold.

As already mentioned above, in this case, size-age classes will correspond to these ways. They can be ranked according to their size, which determines the intensiveness of consumption of natural forage resources per individual.

In this case, to study of the role of biodiversity for different regimes of the system’s usage of resources, it seems reasonable to compare RMUR and RLUR by the character of the distribution of combinations of the values of fish body’s variability. In this paper, we are dealing with the following parameters:

-mode (Mo) — the parameter indicating the most often occurring value (small, large or medium) of the body weight;

-mode amplitude (Amo) — the parameter indicating the part of observations when Mo occurs;

-the difference between the maximum and minimum body mass observed (Max-Min);

-the maximum of the body weight observed (Max);

Tables 3 and 4 show the idealized trajectories of the system calculated for RLUR and RMUR, respectively, reflecting the set of combinations of Mo, Amo, Max-Min, Max. This set of combinations, as mentioned above, in this case, can be interpreted as a set of the system’s strategies of a fish population for using the resource of a natural forage reserve at different levels of its limitation and accessibility. Time moments in the ITS correspond to strategies-combinations. Comparison of the appearance of ITSs, presented in Table 3 and Table 4, enables to find sufficiently expressive differences between RLUR and RMUR in the character of relationships between Mo and Amo. We keep in mind the following. In Table 3, there are three time moments (1st, 2nd, and 3d), for which the maximum (three scores) of Mo and Amo values are observed simultaneously. This systemic effect can have the following biological meaning. Restriction of the resources of a natural forage reserve, which is an attribute of RLUR, leads to increasing the level of food competition between different size-age classes. Combinations-strategies of the coincidence of high values of Mo and Amo suggest that the dominant largest size-age class, as a result of food competition, forces out other classes. Due to this, the proportion of such the class takes relatively high values in the population. This aspect is absent or much less important in RMUR, and respectively, in the ITS presented in Table 4. There are no time moments for which we observe high values of Mo and Amo simultaneously. Thus, the systemic effect associated with the ratio of the values of Mo and Amo in combinations-strategies can be a diagnostic feature that makes it possible to distinguish between RMUR and RLUR. Different values of Mo and Amo can be considered as different intensities of different ways to increase the efficiency of usage of natural forage resources: by increasing the body weight of individuals of the dominant size-age class or by increasing the proportion of individuals of this class in the population. Different degrees of evenness of intensiveness of usage of mentioned ways correspond to different combinations of values of Mo and Amo.

**Table 3.**
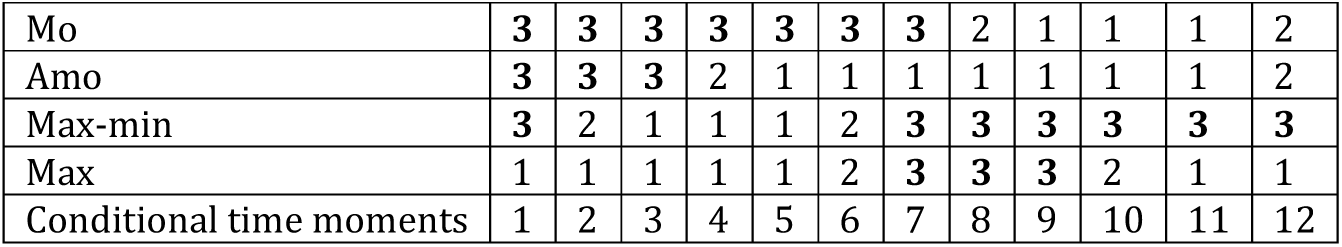
The idealized trajectory of the system representing the dynamics of the values of the parameters of body mass variability of *Karassius karassius* in the period before the experimental population decline.

|  |  |  |  |  |  |  |  |  |  |  |  |  |
| --- | --- | --- | --- | --- | --- | --- | --- | --- | --- | --- | --- | --- |
| Mo | **3** | **3** | **3** | **3** | **3** | **3** | **3** | 2 | 1 | 1 | 1 | 2 |
| Amo | **3** | **3** | **3** | 2 | 1 | 1 | 1 | 1 | 1 | 1 | 1 | 2 |
| Max-min | **3** | 2 | 1 | 1 | 1 | 2 | **3** | **3** | **3** | **3** | **3** | **3** |
| Max | 1 | 1 | 1 | 1 | 1 | 2 | **3** | **3** | **3** | 2 | 1 | 1 |
| Conditional time moments | 1 | 2 | 3 | 4 | 5 | 6 | 7 | 8 | 9 | 10 | 11 | 12 |

**Table 4.** Idealized trajectory of the system, representing the dynamics of the values of the parameters of body mass variability of *Karassius karassius* in the period after the experimental population decline. Notation as in Table 3.

|  |  |  |  |  |  |  |  |  |  |  |  |
| --- | --- | --- | --- | --- | --- | --- | --- | --- | --- | --- | --- |
| Mo | 1 | 1 | 1 | 1 | 1 | 1 | 2 | 3 | 3 | 3 | 2 |
| Amo | 1 | 2 | 2 | 1 | 1 | 1 | 1 | 1 | 1 | 1 | 1 |
| Max-min | 1 | 2 | 3 | 3 | 3 | 3 | 2 | 1 | 1 | 1 | 1 |
| Max | 1 | 1 | 1 | 1 | 2 | 3 | 3 | 3 | 2 | 1 | 1 |
| Conditional time moments | 1 | 2 | 3 | 4 | 5 | 6 | 7 | 8 | 9 | 10 | 11 |

Rows – the values of the system’s components — parameters of body mass variability, data in conditional scores (1 — low, 2 — medium, 3 — high). Columns are time moments. The abbreviations of the parameters of body mass variability are explained in the text. The maximum values for the table are indicated in bold.

As we see, the approach to the diagnosis of RMUR and RLUR based on the determining the degree of evenness of different ways for using resources previously proposed and preliminarily grounded in the work [27], devoted to use of the DMDS for a formalized description of the performance of the human cardiovascular system, can be used for study of the behavior of systems at a completely different level of organization of living matter. This confirms the above-suggested working hypothesis.

The conclusion about the prospects of use of this approach along with the DMDS for studying the role of certain aspects of biodiversity in different regimes of resources’ usage by a wide range of biological systems, taking into account the above, seems to be sufficiently sound.

